# Arabidopsis *HIPP27* is a host susceptibility gene for the beet cyst nematode *Heterodera schachtii*

**DOI:** 10.1101/208132

**Authors:** Zoran S. Radakovic, Muhammad Shahzad Anjam, Elizabeth Escobar, Divykriti Chopra, Javier Cabrera, Ana Cláudia Silva, Carolina Escobar, Miroslaw Sobczak, Florian M. W. Grundler, Shahid Siddique

## Abstract

**Summary:** Sedentary plant-parasitic cyst nematodes are obligate biotrophs that infect the roots of their host plant. Their parasitism is based on modification of root cells to form a hypermetabolic syncytium from which the nematodes draw their nutrients. The aim of this study was to identify nematode susceptibility genes in *Arabidopsis thaliana* and to characterize their roles in supporting the parasitism of *Heterodera schachtii*. By selecting genes that were most strongly upregulated in response to cyst nematode infection, we identified HIPP27 (<u>H</u>EAVY METAL-ASSOCIATED ISOPRENYLATED <u>P</u>LANT <u>P</u>ROTEIN 27) as a host susceptibility factor required for beet cyst nematode infection and development. Detailed expression analysis revealed that HIPP27 is a cytoplasmic protein and that *HIPP27* is strongly expressed in leaves, young roots and nematode-induced syncytia. Loss-of-function Arabidopsis *hipp27* mutants exhibited severely reduced susceptibility to *H. schachtii* and abnormal starch accumulation in syncytial and peridermal plastids. Our results suggest that *HIPP27* is a susceptibility gene in Arabidopsis whose loss-of-function reduces plant susceptibility to cyst nematode infection without increasing susceptibility to other pathogens or negatively affecting plant phenotype.

## Introduction

Cyst nematodes are obligate biotrophs that cause extensive yield losses in almost all economically important crops (Jones *et al*., 2013). The lifecycle of the cyst nematode begins when an infective-stage juvenile invades the root, preferentially in the elongation zone. Once inside the root, the juvenile beet cyst nematode travels through various tissue layers until it reaches the vascular cylinder, where it selects a single cell that will become the initial syncytial cell (ISC; Wyss & Zunke, 1986). Upon selection of the ISC, the nematode becomes immobile and releases proteinaceous and non-proteinaceous secretions inside the ISC to promote the formation and function of the syncytium (Hewezi *et al.*, 2015, Gardner *et al.*, 2015, Siddique *et al.*, 2015, Habash *et al.*, 2017). The syncytium is a metabolic sink and serves as the nematode’s only source of nutrients for the remainder of its lifecycle. A plethora of metabolic, proteomic and transcriptomic changes accompany the development of the syncytium (Szakasits *et al*., 2009, Hofmann *et al*., 2010, Elashry *et al*., 2013; Hütten *et al.*, 2015, Siddique *et al.*, 2014a). As the syncytium expands, the nematode differentiates into either a male or female (Trudgill, 1967). Although the mechanism of sex determination is not yet fully understood, host factors such as defence activation and nutrient availability influence the sexual outcomes of cyst nematodes (Siddique *et al*., 2014b, Siddique *et al*., 2015, Mendy *et al.*, 2017, Shah *et al.*, 2017). When there is a surplus of nutrients, more females develop. However, most nematodes develop into males under stress conditions, such as those in resistant plants.

A previous transcriptome analysis showed that expression of a HEAVY METAL-ASSOCIATED ISOPRENYLATED PLANT PROTEIN (HIPP) family gene, *HIPP27*, is strongly upregulated in syncytia induced by the beet cyst nematode *Heterodera schachtii* in Arabidopsis roots (Szakasits *et al*., 2009). Both prokaryotes and eukaryotes have evolved mechanisms to maintain efficient metal homeostasis inside the cell, including the presence of numerous metal transport proteins known as metallochaperones. Most metallochaperones contain a heavy metal-binding domain (HMA; HMA, pfam00403.6) with a highly conserved CysXXCys motif and a βαββαβ-fold structure for binding Cu^+^, Cd^2+^ or Zn^2+^ (Tehseen *et al*., 2010). Many different families of metallochaperones function in plant defences against the toxicity of metals such as cadmium, chromium and aluminium. In addition, several plant immune receptors carry HMA domains, indicating that HMA likely plays a role in plant defence against pathogens (Sarris *et al.*, 2016).

In addition to the HMA domain, members of the large HIPP family contain a C-terminal isoprenylation motif. HIPPs are present only in vascular plants and are involved in a variety of biological processes, including heavy metal homeostasis and detoxification (Tehseen *et al*., 2009), transcriptional responses to abiotic stresses such as drought and cold (Barth *et al*., 2009), and plant–pathogen interactions (de Abreu-Neto *et al*., 2013; Zschiesche *et al*., 2015). However, the mechanistic details underlying the roles of HIPPs in these biological processes remain mostly unknown. In *A. thaliana*, HIPPs make up the largest metallochaperone family, comprising 45 members divided into seven distinct classes (Tehseen *et al.*, 2009). The best-characterized member of this family in Arabidopsis is HIPP3, a zinc-binding protein that functions as an upstream regulator of the salicylate-dependent pathway during pathogen infection (Zschiesche *et al*., 2015). As described above, expression of *HIPP27* is strongly upregulated in syncytia induced by *H. schachtii* in Arabidopsis roots. Therefore, we hypothesized that HIPP27 might be an important host susceptibility factor in beet cyst nematode parasitism. In the present study, we investigated the role of HIPP27 in the interaction between *H. schachtii* and Arabidopsis. Our data show that *HIPP27* facilitates cyst nematode infection and the efficient development of nematode feeding sites in Arabidopsis roots.

## Results and Discussion

### Selection of *HIPP27* as a candidate Arabidopsis host susceptibility gene for *H. schachtii*

The main objective of this study was to identify new target genes that could play a role in the susceptibility of Arabidopsis to *H. schachtii*. We mined previously published transcriptomic data and selected the top 100 genes that are the most strongly upregulated in the syncytium compared to uninfected control roots (Szakasits *et al*., 2009). We then performed a literature search, and all genes that play a role in crucial metabolic processes, such as photosynthesis and sugar metabolism, were excluded. This step was aimed at minimizing the risk of selecting for which the loss-of-function mutant may show severe growth phenotypes. We also eliminated all genes whose loss-of-function mutants have been previously shown to exhibit severe root or shoot phenotypes. Next, BLAST searches were used to identify candidate genes for which only a single copy ortholog is present in the genome of the natural host of *H. schachtii* i.e., *Beta vulgaris* (Dohm *et al*., 2014). Using this process, we ultimately selected 10 candidate host susceptibility genes (**Table 1**). The expression of these genes was validated in Arabidopsis plants that were grown *in vitro* and infected with *H. schachtii*. RNA was extracted from hand-dissected root segments containing syncytia that were sampled at 15 dpi and was used to analyse the expression of the candidate genes via qRT-PCR. The results confirmed the upregulation of all 10 candidate genes (**Table 1**), although the intensity of fold change was substantially lower than in the microarray data. This discrepancy between the qPCR and microarray data can likely be attributed to the different syncytial material used in the two cases: whereas microaspirated syncytial cell contents were used for microarray analysis, our qRT-PCR analysis was performed using cut syncytia containing the surrounding non-syncytial root cells, which may have diluted syncytium-specific mRNA expression.

**Table 1:** Validation of 10 selected candidate genes expression in syncytia induced in Arabidopsis roots by beet cyst nematode. For microarrays, data from microaspirated syncytia at 5 and 15 dpi were pooled and compared with control roots (Szakasits *et al*., 2009). For qPCR, values represent relative fold change in infected root segment containing syncytia as compared to control uninfected root. *18S* and *UBP22* were used as housekeeping genes to normalize the data. All values are means of three biological replicates +/-SE.

| Gene | Locus | Fold change compared to uninfected control |  |
| --- | --- | --- | --- |
|  |  | Microarrays (5+15 dpi) | qRT-PCR 15dpi |
| *ENH1* | AT5G17170 | 27.9 | 2.6 +/- 0.26 |
| *UGP2* | AT5G17310 | 29.9 | 2.6 +/- 0.56 |
| -- | AT4G24830 | 29.9 | 3.3 +/- 0.59 |
| *IMD2* | AT1G80560 | 32.0 | 7.3 +/- 4.7 |
| *NPC6* | AT3G48610 | 39.3 | 30.0 +/- 11.6 |
| *AD11A3* | AT2G24270 | 48.5 | 44.5 +/- 16.9 |
| *TRX1* | AT3G51030 | 48.5 | 11.2 +/- 16.9 |
| *CCR2* | AT1G80820 | 520 | 15.47 +/- 3.48 |
| *FAD6* | AT4G30950 | 52.0 | 5.04 +/- 1.71 |
| *HIPP27* | AT5G66110 | 68.6 | 24.81 +/- 2.88 |

We obtained loss-of-function homozygous mutants for the 10 candidate genes and analysed them for beet cyst nematode susceptibility via preliminary pathogenicity assays (**Figure S1**). Among these, a mutant for the candidate gene *HIPP27* showed a particularly significant decrease in susceptibility to beet cyst nematode and was therefore selected for further molecular characterization.

### *HIPP27* is strongly upregulated in the syncytium

To investigate the expression patterns of *HIPP27* in various organs of Arabidopsis plants, we cloned the putative promoter region (472 bp) upstream of the translation start codon of *HIPP27*, producing the *pHIPP27:GUS* construct. Arabidopsis plants were transformed with *pHIPP27:GUS* and three independent homozygous lines were obtained. The general expression pattern of *pHIPP27:GUS* was then assessed in various plant organs. For uninfected lines, GUS staining was detected ubiquitously in all tissues examined, with particularly strong expression in young leaves (**Fig. 1A–C**). In roots, GUS staining was detected in the vascular cylinder of young lateral roots (**Figure 1D**); however, no staining was detected in the elongation zone and in root tips (**Figure 1E**). Plants harbouring *pHIPP27:GUS* were then infected with nematodes and stained for GUS activity at different time-points after infection, i.e., 1, 3, 5 and 10 dpi, to cover the different developmental stages of nematode development (**Fig. 1F**). We detected strong GUS expression at 3 dpi (**Fig. 1G)**, which became intense at 5 dpi (**Fig. 1H**). However, the intensity of GUS staining decreased significantly at 10 dpi (**Fig. 1I**). Notably, GUS expression was generally localized inside the syncytium and extended beyond the syncytium in only a few cases, indicating that the upregulation of *HIPP27* is specific to nematode infection.

**Figure 1:**
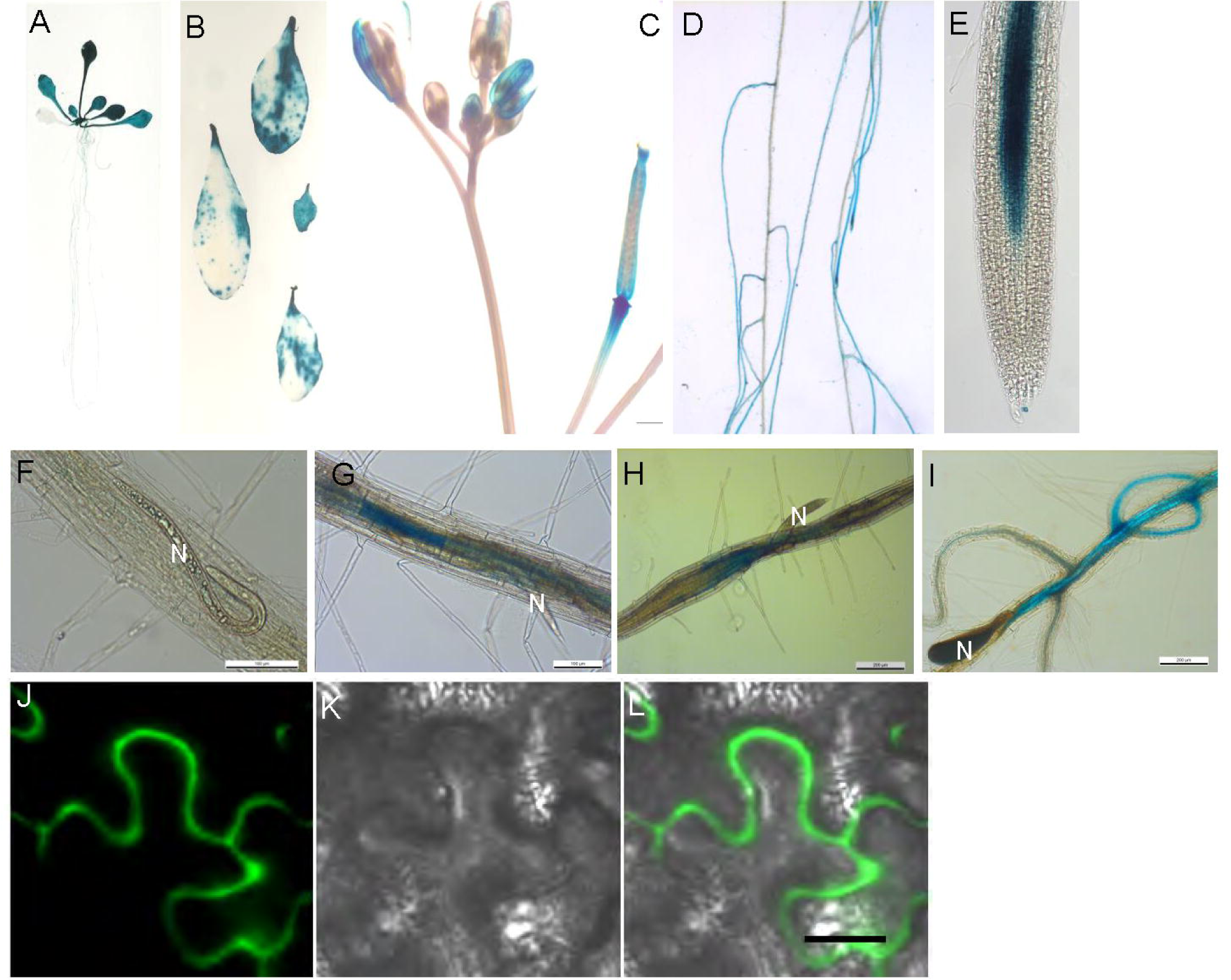
*HIPP27* is strongly upregulated in the syncytium. Expression of *pHIPP27:GUS* in various organs of Arabidopsis. (A) 14-day-old seedling. (B) Leaves from 2–3–week-old plants. (C) Inflorescence. (D and E) 12-day-old root (0 dpi). (F–I) Nematode-infected root segments at 1 (F), 3 (G), 5 (H), and 10 (I) dpi. (J–L) Localization of *35S:HIPP27-GFP* in epidermis of *Nicotiana benthamiana* leaves. N, nematode.

To determine the subcellular localization of HIPP27, HIPP27-GFP was transiently expressed in *Nicotianna benthamiana* leaf epidermis under the control of a constitutive promoter (CaMV 2x35S) and its localization assessed using confocal microscopy. HIPP27-GFP was localized to the cytosolic region of the cell (**Figure 1J–L**). Taken together, these results indicate that HIPP27 is a cytoplasm-localized protein and that *HIPP27* expression is strongly upregulated during early stages of syncytium development. These findings point to the likelihood of importance of this gene in early syncytium development.

### Loss-of-function of *HIPP27* decreases Arabidopsis susceptibility to *H. schachtii*

To further substantiate the role of HIPP27 in the plant’s interaction with cyst nematodes, we obtained loss-of-function T-DNA insertion mutants of *HIPP27* (**Figure S2**). Homozygous lines were selected after genotyping and the lack of *HIPP27* expression in the homozygous mutants was confirmed by RT-PCR (**Figure S3**). To determine whether these *hipp27* mutants showed any impairment in growth and development, we analysed phenotypic parameters in plants grown under sterile and greenhouse conditions. However, we did not observe any significant difference in average dry weight, average flowering time, average root length or average plant height in the mutant compared to the control (**Figure S4A–D**).

To analyse the changes in *hipp27* with regard to nematode susceptibility, we performed infection assays in which several susceptibility parameters were measured. The average number of females, a commonly accepted parameter indicating nematode susceptibility under *in vitro* conditions, was strongly reduced in *hipp27* compared to the control (**Figure 2A**). In addition, the average numbers of males and average total number of invading and developing nematodes were also significantly reduced in *hipp27* compared to Col-0 (**Figure 2A**). Further susceptibility parameters, such as the average sizes of the female and the syncytium, were also significantly reduced in *hipp27* compared to Col-0 (**Figure 2B** and **2C**). For field-grown crops, the most common susceptibility parameters are the average number of eggs per cyst, average size of cysts and the pi/pf value (Final nematode population / Initial nematode population ratio), which represent the reproductive potential of the nematode. Both the average number of eggs per cyst and pi/pf were significantly reduced in *hipp27* compared to Col-0 (**Figure 2D–2G**). Together, these results suggest that HIPP27 plays a vital role in cyst nematode parasitism.

**Figure 2:**
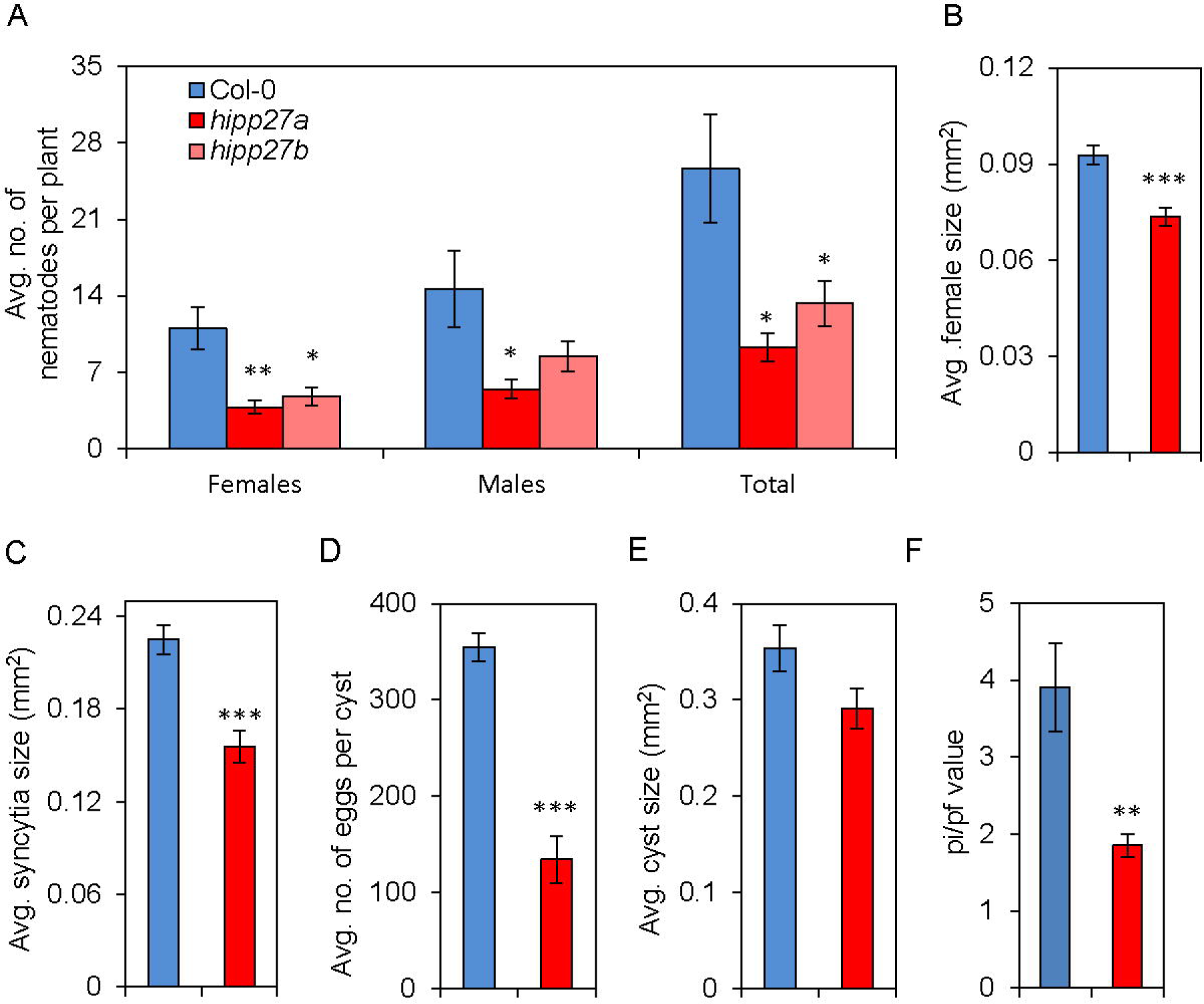
Loss of function of *HIPP27* decreases Arabidopsis susceptibility to *H. schachtii*. (A) Average number of nematodes per plant present in Col-0 and *hipp27* mutant lines at 14 dpi. (B–F) Average sizes of female nematodes (B) and plant syncytia (C), average number of eggs per cyst (D), average cyst sizes (E) and pf/pi values (F), in Col-0 and *hipp27* mutant lines at 14 dpi. (A–G) Bars represent mean ± SE of three independent experiments. Asterisks represent statistically significant difference from corresponding Col-0 value (*t-test*, *P < 0.05, **P < 0.01, ***P < 0.001).

Because loss-of-function of *HIPP27* resulted in a significant decrease in susceptibility to beet cyst nematodes, we hypothesized that overexpressing this gene might increase susceptibility to nematodes. To test this, we produced transgenic lines expressing *HIPP27* under the control of a constitutive promoter in either the Col-0 (*35S:HIPP27*) or *hipp27* (*35S:HIPP27*/*hipp27*) background. We confirmed the increase in transcript abundance in the homozygous lines by qRT-PCR (**Figure 3A**) and performed an infection assay. The average number of female nematodes per plant was significantly higher in the *35S:HIPP27* lines compared to Col-0 (**Figure 3B**). However, other parameters, such as female size and syncytium size, did not differ in these lines compared to the control. By contrast, the *35S:HIPP27*/*hipp27* lines did not show any change in the susceptibility to nematodes compared to the control (**Figure 3C** and **3D**). Altogether, these data suggest that the expression of *HIPP27* is activated upon cyst nematode infection and is required for proper syncytium and nematode development.

**Figure 3:**
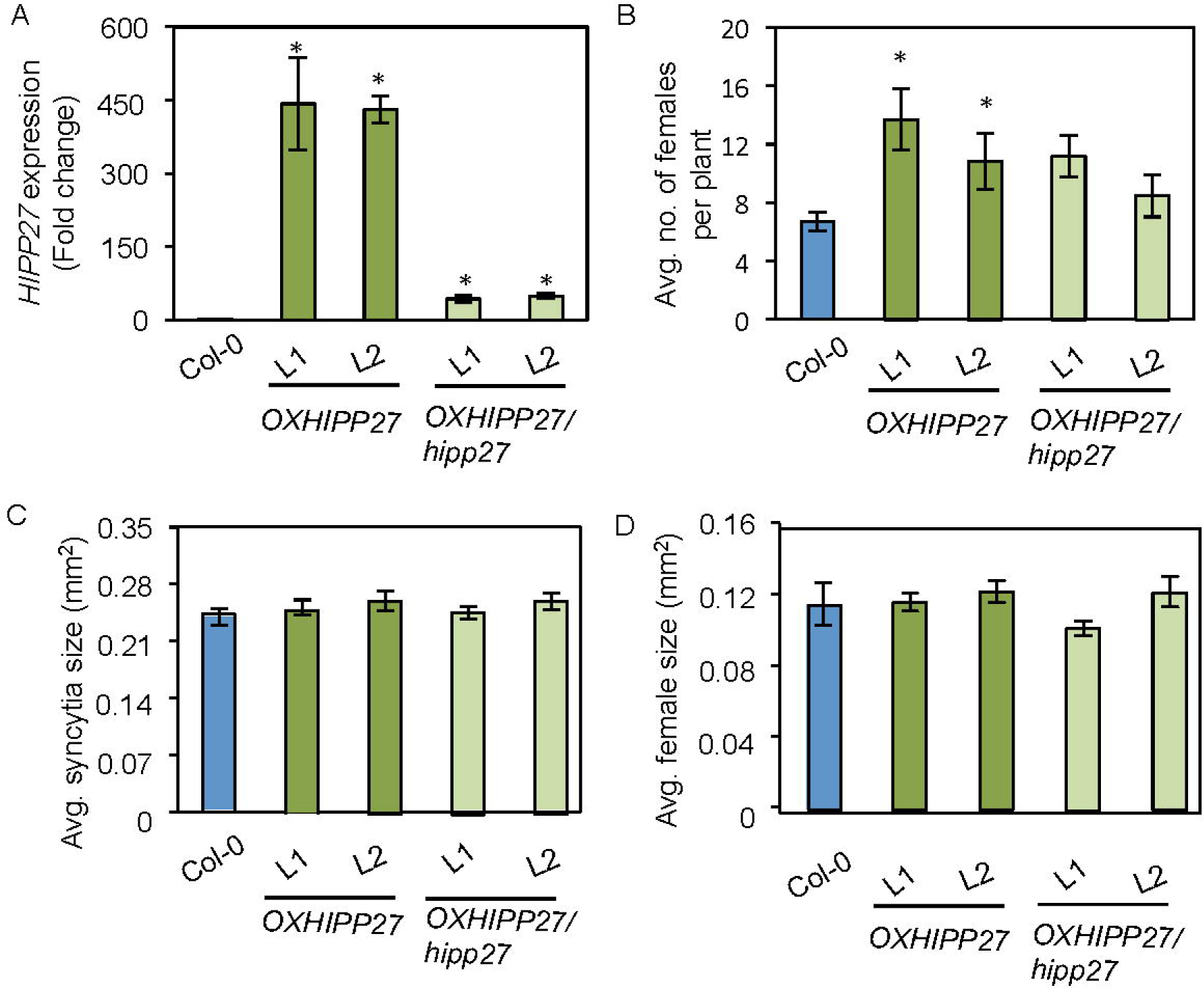
Overexpression of *HIPP27* lead to increased susceptibility to *H. schachtii*. (A) qRT-PCR confirmation of increase of *HIPP27* transcript in overexpression lines of the Col-0 (*OXHIPP27*) or *hipp27* (*OXHIPP27*/*hipp27*) background. (B) Average number of nematodes per plant present in Col-0 and *HIPP27* overexpression lines at 14 dpi. (C and D) Average sizes of female nematodes (C) and syncytia (D) in Col-0 and *HIPP27* overexpression lines at 14 dpi. (A– D) Bars represent mean ± SE of three independent experiments. Asterisks denote significant difference from corresponding Col-0 value (*t-test*, *P < 0.05, **P < 0.01, ***P < 0.001).

### Loss-of-function of *HIPP27* does not lead to increased susceptibility to other pathogens

We wanted to determine whether the *hipp27* mutant exhibits increased susceptibility to other pathogens. Therefore, we performed infection assays using the gall-forming nematode *Meloidogyne javanica* and the necrotrophic fungus *B. cinerea*. We did not observe differences between the wild type and mutant in the average size of necrotic lesions caused by *Botrytis cinerea* (**Figure 4A**). In addition, although the number of galls formed by *M. javanica* in *hipp27* decreased slightly as compared to the Col-0 controls, the difference was not significant (**Figure 4B**). To further rule out a putative role for HIPP27 during gall formation, we compared the diameters of the galls in the wild-type and *hipp27* mutant lines, finding no significant differences between them (**Figure 4C**). Moreover, the reproduction of the RKNs, measured as the number of egg masses per gall, was statistically unaffected in *hipp27* (**Figure 4D**). Together, these data pointed to a cyst nematode-specific role of *HIPP27* in supporting parasitism.

**Figure 4:**
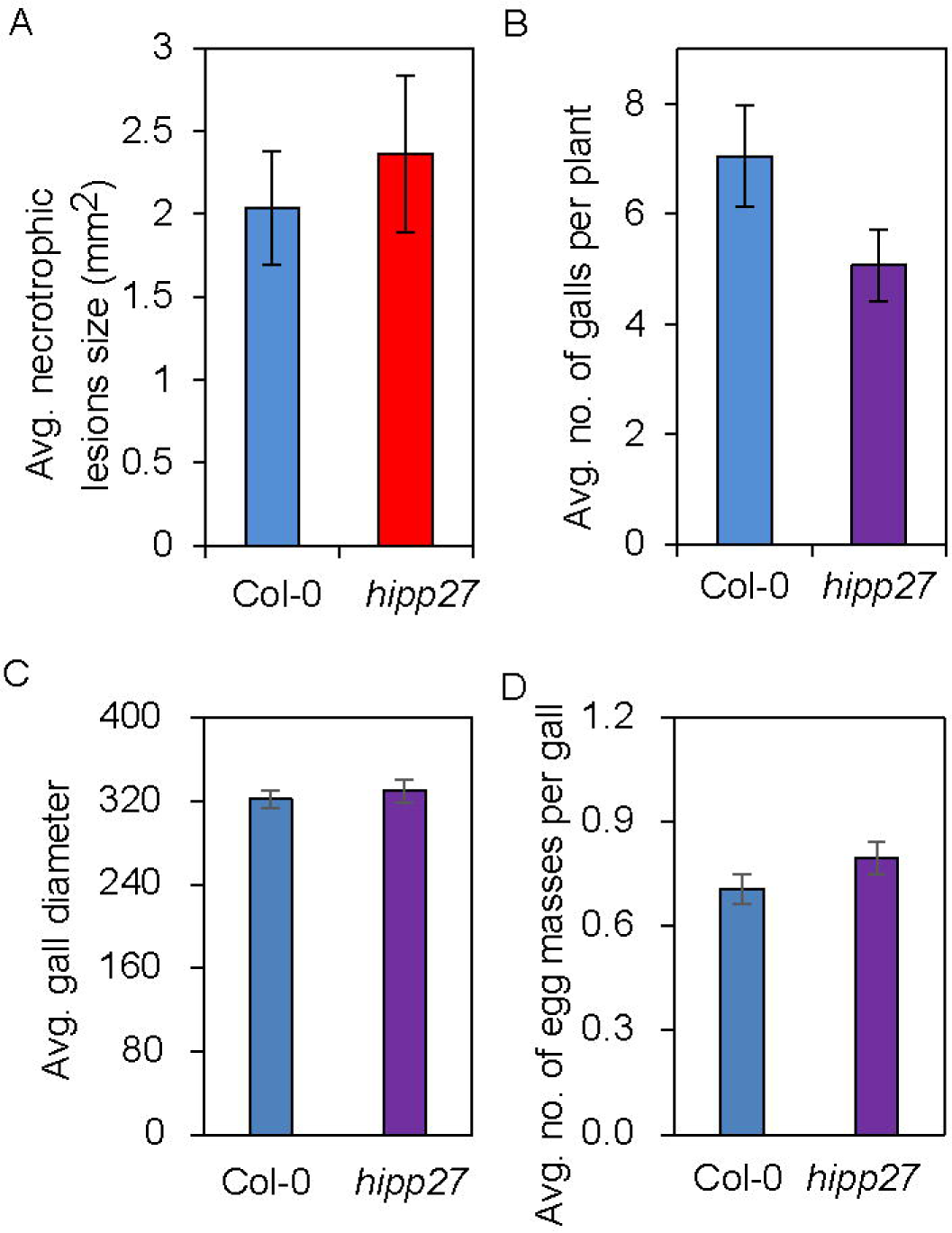
Loss of function of *HIPP27* does not alter susceptibility to other pathogens. (A) Average size of lesions induced by *B. cinerea* in Col-0 and *hipp27* lines. (B) Average number of galls per plant in Col-0 and *hipp27* mutant at 21 dpi with *M. javanica*. (C) Gall size calculated as the diameter of galls from Col-0 and *hipp27* at 14 dpi with *M. javanica*. (D) Number of egg masses formed per gall formed by *M. javanica* in Col-0 and *hipp27* two months after infection. Statistical analysis was performed with three independent experiments using a Student’s *t*-test (no significant differences from Col-0 were observed). Bars represent mean ± SE of three independent experiments.

### Loss of function of *HIPP27* does not impair plant basal defence

To investigate whether the reduced susceptibility of *hipp27* plants to *H. schachtii* is due to altered plant defence responses, we measured the ROS levels induced over a 60-minute period upon treatment of leaves with a well-known immunopeptide, flg22. We did not observe significant changes in ROS levels (**Figures 5A** and **5B**), indicating that the *hipp27* mutant does not have impaired plant basal defence responses. To further confirm this finding, we performed qRT-PCR analysis to examine the expression of a few plant basal defence marker genes in uninfected shoots: *PR1* and *PR2* are induced by salicylic acid, *PDF1.2* and *HEL* are marker genes for jasmonic acid signalling, and *ERF6* and *ACS2* are involved in ethylene signalling. The expression levels of all marker genes in the *hipp27* mutant were similar to those of Col-0 (**Figure 5C**). These results reinforce the conclusion that *HIPP27* has a specific role in the interaction with cyst nematodes, as an increase in the plant basal defence would lead to a resistance to other pathogens (**Figure 4A–D**).

**Figure 5:**
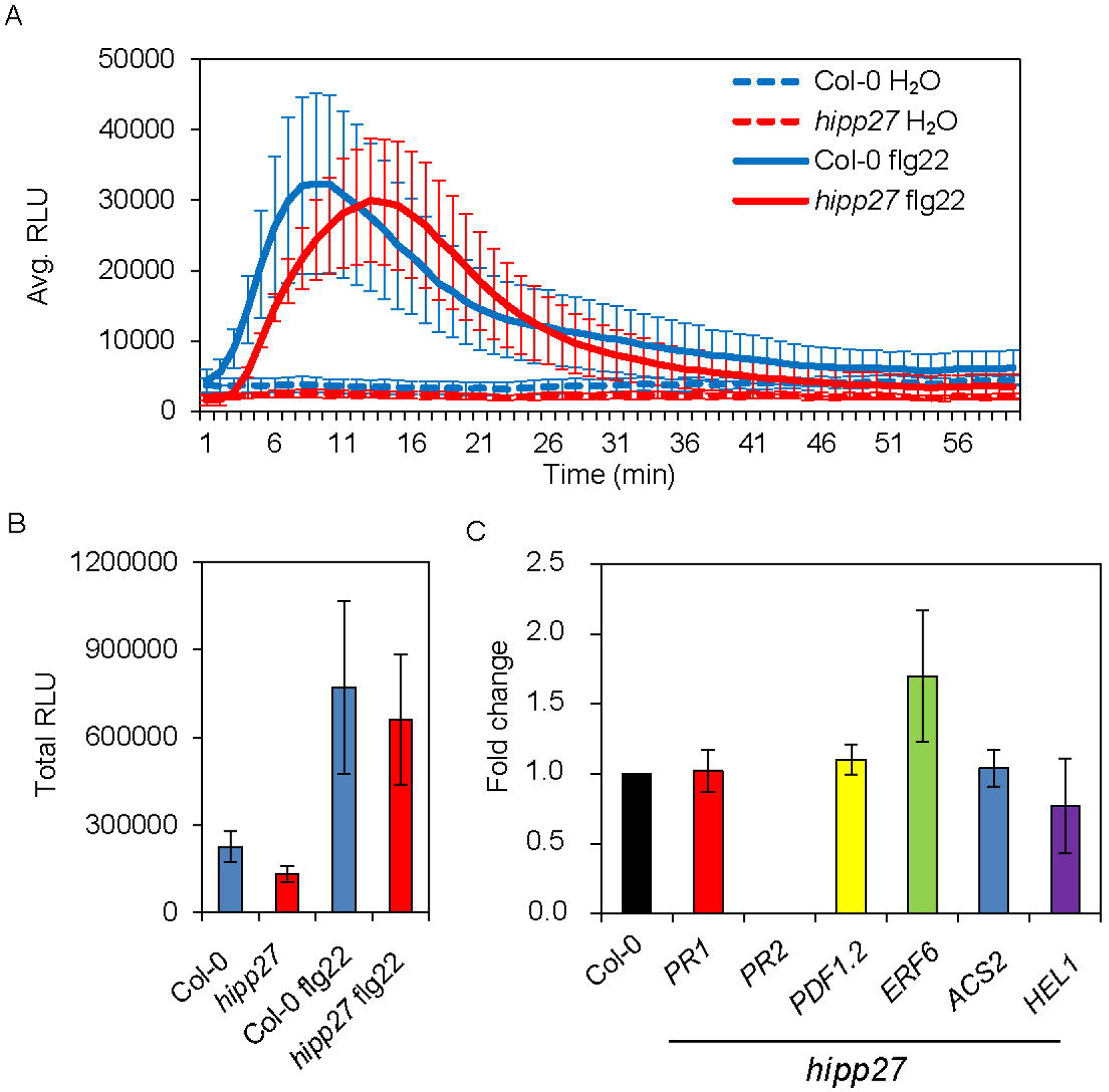
Loss of function of *HIPP27* does not impair plant basal defence. (A) ROS burst induced by flg22 during 60 min of measurement in Col-0 and *hipp27* mutant expressed in relative light units (RLU). (B) Total RLU ROS during 60 minutes of exposure to flg22 in Col-0 and *hipp27* mutant. (C) qRT-PCR gene expression analysis of defence-related genes in uninfected shoots of Col-0 and *hipp27* mutant. Statistical analysis was performed with three independent experiments per line using a Student’s *t*-test (no significant differences from Col-0 were observed). Bars represent mean ± SE of three independent experiments.

### Loss of function of *HIPP27* causes physiological or metabolic abnormalities

Finally, we examined the anatomical and ultrastructural organisation of syncytia induced by the beet cyst nematode in roots of wild-type Col-0 and *hipp27* mutant plants (**Figure 6**). Comparison of light microscopy images of sections taken through similar regions of nematode-induced syncytia showed no apparent differences between the syncytia induced in the two genotypes (**Figure 6A–D** versus **6E-H**). Next to the nematode head, the syncytia were relatively small on cross sections and cells not incorporated into them divided at different planes, forming groups of neoplastic cells (**Figure 6A, 6C, 6E** and **6G**) similar to those described by Sobczak *et al*. (1997). The extent of destruction around the nematode head was also similar in both plant genotypes (**Figure 6A, 6C, 6E** and **6G**). Sections from the middle and the widest region of syncytia (distant from the nematode head) showed that if the syncytium had been induced in young roots without fully differentiated primary xylem, it occupied the central position inside the vascular cylinder and it became surrounded by regularly dividing pericyclic cells forming periderm, a secondary cover tissue (**Figure 6B** and **6F**). However, if the syncytium had been induced in root region where primary xylem was fully differentiated, it developed the primary xylem on only one side whereas secondary xylem elements differentiated on the opposite side (**Figure 6D** and **6G**). In both cases, the pericyclic cells divided and formed periderm around the entire vascular cylinder (**Figure 6B, 6D, 6F** and **6H**). No visible differences in the extent of syncytial element hypertrophy or the number and size of cell wall openings were evident, indicating that the anatomical structure of syncytia induced in *hipp27* mutant was undisturbed (**Figure 6A–H**) although their dimensions were smaller compared to Col-0 (**Figure 2**).

**Figure 6:**
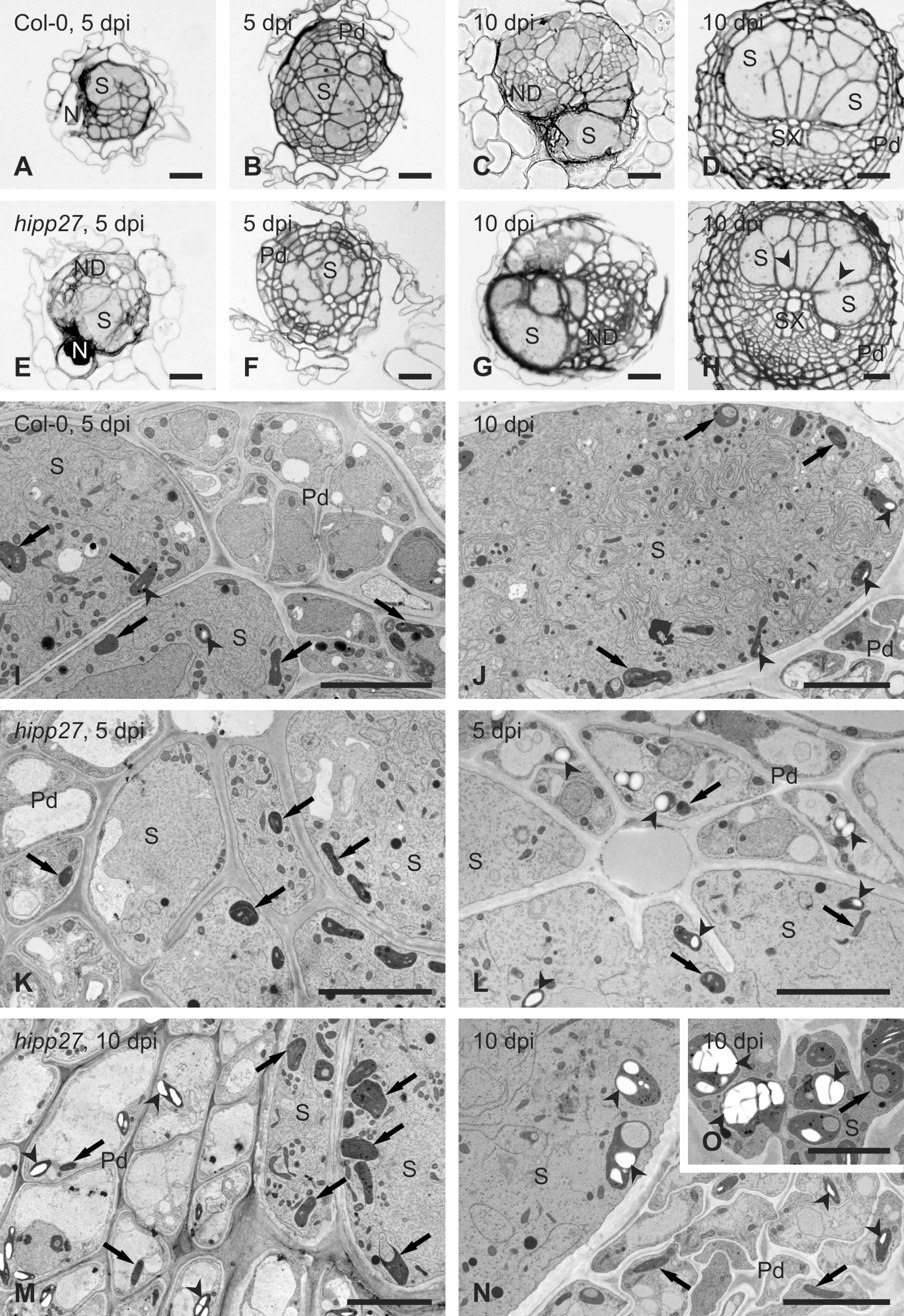
Loss of function of *HIPP27* causes physiological or metabolic abnormalities. Light (A-H) and transmission electron microscopy (I-O) images of cross sections of syncytia at 5 (A, B, E, F, I, K and L), and 10 dpi (C, D, G, H, J and M-O) induced in roots of Col-0 wild-type plants (A-D, I and J) and *hipp27* mutant (E-H and K-O). Light microscopy images were taken close to the nematode head (A, C, E and G) or through the widest regions of syncytia (B, D, E and H). Arrowheads indicate selected starch grains in plastids, arrows point to plastids. *Abbreviations*: N, nematode; ND, neoplastic divisions; Pd, periderm; S, syncytium; SX, secondary xylem. Scale bars: 20 µm (A-H), 5 µm (I-O).

At ultrastructural level, syncytia induced in Col-0 and *hipp27* plants were similar at 5 dpi (**Figure 6I** and **6K**). Their cytoplasm proliferated and became electron dense, with numbers of plastids, mitochondria and endoplasmic reticulum structures increasing, whereas central vacuoles disappeared. Only few syncytial plastids contained very small starch grains. However, serial sectioning of syncytia showed that syncytia induced in *hipp27* roots often differed in their ultrastructure along their axis. In the region close to the nematode head and at the leading edge (the most remote part of the syncytium), syncytia had typical ultrastructural organisation (**Figure 6K**). By contrast, in the middle region of the syncytium, the cytoplasm was relatively electron translucent, endoplasmic reticulum structures were weakly developed, and large starch grains were formed in numerous syncytial plastids and plastids of peridermal cells next to the syncytium (**Figure 6L**). In samples collected at 10 dpi, syncytial protoplasts had typical ultrastructure in syncytia induced in wild-type Col-0 plants (**Figure 6J**). Here again, small starch grains were present in only a few syncytial plastids and they were absent in cells surrounding the syncytium. By contrast, starch grains were frequently present in peridermal plastids next to syncytia induced in *hipp27* roots (**Figure 6M** and **6N**). Inside syncytia, plastids present in regions with well-developed endoplasmic reticulum usually did not contain starch grains (**Figure 6M**), whereas plastids present in regions with electron translucent cytoplasm and poorly developed endoplasmic reticulum usually contained large starch grains (**Figure 6N** and **6O**). Some of these plastids acquired enormously large sizes (**Figure 6O**) and could be clearly recognised even on anatomical sections (**Figure 6H**). Taken together, our ultrastructural analysis showed that loss of function of HIPP27 leads to physiological and metabolic abnormalities including accumulation of large starch grains.

### Conclusion

This study represents the first investigation of the role of an HIPP family protein in plant– nematode interactions. We showed that the expression of *HIPP27* is specifically and highly upregulated in syncytia induced by *H. schachtii* in Arabidopsis roots. The strong GUS expression driven by the *HIPP27* promoter inside the syncytium points to the involvement of *HIPP27* in the establishment and development of syncytia. Indeed, analysis of loss-of-function mutants and overexpression lines showed that *HIPP27* expression supports infection of the host roots by the beet cyst nematode. In contrast to its expression in cyst nematode-induced syncytia, *HIPP27* expression was not differentially regulated in feeding sites induced by root-knot nematodes (Cabrera *et al*., 2014). These observations indicate that the role of *HIPP27* in facilitating parasitism is restricted to that of cyst nematodes. This hypothesis is further supported by results from pathogenicity assays with *M. javanica* and *B. cinerea*, in which we did not observe any significant difference in susceptibility between Col-0 and the *hipp27* mutant.

HIPP27 belongs to Cluster IV of the HIPP family in Arabidopsis, which is notable for the presence of a conserved Asp residue preceding the metal binding motif (CysXXCys). Notably, the expression of *HIPP27* in yeast confers slight increase of Cd resistance to a Cd-sensitive yeast strain, likely due to binding of Cd by HIPP27 in the cytosol (Tehseen *et al*., 2010). Several HIPP family members including HIPP26 and HIPP27 have been shown to interact via the HMA domain with the drought stress-related zinc finger transcription factor 29 (ATHB29) and UBIQUITIN-SPECIFIC PROTEASE 16 (UBP16) (Barth *et al*., 2009, Zhao *et al*., 2013). In addition, *hipp26* mutants display altered expression of ATHB29-regulated dehydration response genes compared to wild type (Barth *et al*., 2009). Whether HIPP27 plays a similar role in ATHB29-mediated gene regulation remains to be explored. Previous microarray analysis using microaspirated *H. schachtii* syncytium protoplasts show that the expression of both *ATHB29* and *UBP16* is downregulated in syncytia as compared to uninfected control roots. By contrast, *HIPP27* is highly upregulated in syncytia (Szakasits *et al*., 2009). These observations make it likely that HIPP27 acts independently of ATHB29 to regulate infection by *H. schachtii*.

Our data showed no difference in the activation of the ROS burst or in the expression of defence genes between Col-0 and *hipp27* mutant lines. Based on these results, we propose that expression of *HIPP27* is required for maintaining the optimal development or functioning of the syncytium. This hypothesis is supported by the observation that the syncytium size was significantly reduced in *hipp27* compared to Col-0. The syncytium is a strong metabolic sink that serves as the only source of nutrients for developing nematodes, and the maintenance of metal homeostasis in the syncytium must be tightly regulated. Considering the role of HIPP family members as metallochaperones, it is plausible that HIPP27 functions in metal transport and homeostasis in the syncytium. Indeed, microscopic observations confirmed that lack of HIPP27 protein does not influence the general developmental pattern of syncytia, but causes physiological or metabolic disorders leading to the accumulation of phloem-provided saccharides as starch grains in peridermal and syncytial plastids. These sugars may become unavailable to parasitic juveniles, leading to their disturbed development. Hofmann *et al*. (2008) have shown that the number of starch grains increases in syncytia during nematode moult when no food is taken up by the associated juvenile, as well as in degraded syncytia of adult males and prematurely degraded syncytia associated with females. As no clearly degraded syncytia were found in our experiments, it can be speculated that starch accumulation and its potential unavailability for the nematodes is the very early feature of syncytium degradation and that this process impairs the development of beet cyst nematodes on *hipp27* roots. However, further work is required to elucidate the precise role of HIPP27 in cyst nematode parasitism.

In conclusion, we identified *HIPP27* as a host susceptibility gene whose deletion reduces plant susceptibility to cyst nematodes without increasing susceptibility to other pathogens. The lack of developmental phenotypes in *hipp27* plants highlights the potential of using *HIPP27* in the breeding of nematode-resistant crop plants.

## Experimental Procedures

### Plant growth and beet cyst nematode infection

*Arabidopsis thaliana* seeds were sterilized for 5 min with 1.2% NaClO, followed by three consecutive washes with autoclaved ddH_2_O. Sterilized seeds were sown in KNOP medium, and 12-day-old plants were infected with 70–80 surface-sterilized J2 individuals. For surface sterilization, hatched J2s were washed with 0.05% HgCl_2_, followed by three consecutive washes with autoclaved ddH_2_O (Sijmons *et al.*, 1991, Siddique *et al*., 2015). Two weeks after inoculation, nematode susceptibility parameters such as average number of females, average number of males, and average number of total nematodes per plant were quantified under a Leica S4E stereo microscope (Leica Microsystems, Germany), and average female and syncytium sizes were measured under a Leica M 165C stereo microscope (Leica Microsystems) equipped with Leica LAS v4.3 image analysis software (Leica Microsystems). Two months later, cyst size and the average number of eggs per cyst were measured and counted, respectively. The pi/pf values were calculated by collecting all developed cysts, crushing them in 1 ml 6% (w/v) NaClO (to avoid egg clustering) and counting the number of eggs per cyst.

### Infection assays *Meloidogyne javanica* and *Botrytis cinerea*

Infection assays using *M. javanica* were performed as described previously (Cabrera *et al*., 2014). In brief, Arabidopsis plants were grown aseptically on 0.3% Gamborg’s medium (Gamborg *et al*., 1968) supplemented with 1.5% (w/v) sucrose for 14 days. Fourteen-days-old plants were inoculated with 70–100 *M. javanica* J2s per plant. The infected plants were incubated at 23°C and 16h:8h, light:dark photoperiod. The number of galls was counted at 21 dpi. Gall diameters were measured at 14 dpi using ImageJ. The infection assays with *B. cinerea* were performed as described previously (Lozano-Torres *et al*., 2014). In brief, 5 µl drops of fungal spores at a concentration of 5x10^5^ were inoculated onto four-week-old Arabidopsis plants grown in soil under greenhouse conditions. The infected plants were incubated for three days in the dark at 20°C and 100% relative humidity. Plant susceptibility was estimated by measuring the sizes of necrotic areas at 3 dpi in the same manner as used to measure the sizes of syncytia or females (Siddique *et al*., 2014b).

### Genotyping and expression analysis of the mutants

Arabidopsis SAIL lines (*hipp27a*, SAIL_167_B06; *hipp27b*, SAIL_675_E09) were ordered from NASC stock centre. Homozygosity of the lines was tested by extracting plant DNA using the CTAB method, after which PCR with SAIL primers was conducted to confirm the T-DNA insertions in these lines. PCR was performed on a C100^TM^ Thermal Cycle (Bio-Rad Laboratories, USA). Visual observation of homozygosity and expression analysis were performed using Gel Doc^TM^ (Bio-Rad Laboratories, USA), together with Lab 3.0 software. The primer sequences are listed in Supplementary Table 1.

### qRT-PCR gene expression analysis

To measure transcript abundance, RNA from mutant leaves was extracted using Nucleospin RNA XS (Macherey-Nagel, Germany) according to the manufacturer’s protocol. The RNA concentration was measured with a NanoDrop (Thermo Fisher Scientific, USA), and cDNA was prepared using a High Capacity cDNA Reverse Transcript kit (Life Technologies, UK). Transcript abundance was measured on a StepOnePlus Real-Time PCR System (Applied Biosynthesis, USA) as described by Pfaffl (Pfaffl, 2001); *18S* and *UBP22* were used as internal controls.

### Plant phenotyping

Phenotypic parameters of Arabidopsis Col-0 and *hipp27* lines were measured using soil-grown plants in a greenhouse and plants grown in Petri dishes on KNOP medium under sterile conditions. Measurements were conducted under a Leica M 165C stereo microscope (Leica Microsystems) equipped with a Leica LAS v4.3 image analysis software (Leica Microsystems) as previously described or manually using a ruler and balance following standard phenotyping protocols (Bolle, 2009).

### Gateway cloning and plant transformation

Gateway cloning was used to clone the *HIPP27* promoter (472 bp upstream of *HIPP27*) and to clone the *HIPP27* CDS into the pDONR207 vector. Briefly, primers with attB extensions were designed and the PCR product was inserted into the pDONR207 vector using Gateway^®^ BP Clonase^TM^ II Enzyme mix (Invitrogen). The pEntry207 vector was transformed into *Escherichia. coli* (DH5α) competent cells using the heat shock method. The pEntry207 vector was then extracted using a Nucleospin^®^ plasmid extraction kit (Machinery-Nagel, Germany), and homologous recombination of the gene or promoter into the destination vector was performed using Gateway^®^ LR Clonase^TM^ II Enzyme mix (Invitrogen, USA). For *HIPP27:GUS*, the destination vector was pMDC162, for *HIPP27:GFP*, the destination vector was pMDC107 and for *35S:HIPP27*, the destination vector was pMDC32 (Curtis & Grossniklaus, 2003). Arabidopsis plants were transformed via the floral-dip method (Clough & Bent, 1998).

### GUS staining

The *HIPP27:GUS* reporter line was introgressed into the Col-0 background, and homozygous lines for the reporter gene were selected in subsequent generations using selection marker genes and genotyping. Histochemical GUS analysis was performed according to Siddique *et al*. (2009). Briefly, the *HIPP27:GUS* lines were grown in Knop medium as described above. The roots were submerged in X-Gluc for 6 hours at 37°C at 1, 3, 5 and 10 dpi. The number of stained syncytia was counted, and photographs were taken under a DMI 4000B microscope (Leica, Microsystems).

### *Nicotiana benthamiana* infiltration assay

The coding region of *HIPP27* without the stop codon was amplified using Gateway forward and reverse primers as given in **Table S1**. The amplified product was cloned into pDONR207 using Gateway^®^ BP Clonase^TM^ II Enzyme mix (Invitrogen). The gene was shuttled into destination vector i.e. pMDC83 using Gateway^®^ LR Clonase^TM^ II Enzyme mix (Invitrogen). The *HIPP27:GFP* construct was transiently expressed in the epidermis of *Nicotianna benthamiana* leaves under the control of the 35S promoter. The GFP fluorescence in 6-week-old *N. benthamiana* leaves was observed under an LSM 710 confocal microscope (Carl Zeiss, Germany).

### Oxidative burst assay

ROS production was measured on a 96-well illuminometer (Mithras LB 940; Berthold Technologies) according to Mendy *et al*. (2017). Briefly, leaf disks 0.5 cm in diameter were cut from 12-day-old Arabidopsis plants and incubated in ddH_2_O for 12 h in the dark. Each leaf disk was placed into the well of a 96-well plate containing 15 µl of 20 µg ml^-1^ horseradish peroxidase and 35 µl of 10 mM 8-amino-5chloro-2,3-dihydro-7phenyl(3,4-d) pyridazine sodium salt (L-012, Wako Chemicals). ROS bursts were induced with 50 µl flg22 (100 µM), with ddH_2_O used as a negative control. Light emission was measured for 60 min, and results were obtained using the software supplied with the instrument.

### Anatomical and ultrastructural analyses

Root segments containing syncytia were dissected from *in vitro*-grown and *H. schachtii* inoculated Arabidopsis Col-0 and *hipp27a* plants at 5 and 10 dpi. They were chemically fixed in a mixture of aldehydes and processed for microscopic examinations as described by Golinowski *et al*. (1996) and Sobczak *et al*. (1997). After embedding in epoxy resin, the samples were serially sectioned for light microscopy examinations at selected places ultrathin sections were taken for transmission electron microscopy (Siddique *et al*., 2014b). Obtained images were cropped, resized and adjusted for similar contrast and brightness using Adobe Photoshop software.

### Bioinformatics analysis

To select the 100 most highly upregulated genes in the 5+15 dpi syncytium, the NEMATIC program was used (Cabrera *et al.*, 2014). Orthologs of Arabidopsis genes were identified using The *Beta vulgaris* Resource (http://bvseq.molgen.mpg.de/index.shtml); Arabidopsis genes sharing at least 60% amino acid sequence identity with sugar beet and without additional homologs were selected. Primers were designed using primer3 online software (http://bioinfo.ut.ee/primer3-0.4.0/). SAIL primers were obtained from the SALK website (http://signal.salk.edu/tdnaprimers.2.html). A list of all primers is presented in **Table S1**.

### Statistical data analysis

Results are presented as a mean ± SE from at least three independent experiments. Statistically significant differences were defined using Microsoft Office Excel and extension XL Statistics with Student’s *t*-test (p<0.05).

## Acknowledgements

This work was funded through the financial support of the Federal Ministry of Education and Research (BMBF), Germany (PLANT-KBBE; Project number, 031A326B). This work was also supported by the Spanish government (grants AGL2013-48787; AGL2016-75287-R to CE). ACS was supported by a fellowship from the University of Castilla-La Mancha, co-funded by the European Social Fund. JC is supported by a Cytema-Santander contract from UCLM. We are grateful to Samer Habash and Badou Mendy for help with many experiments and to Stephan Neumann and Gisela Sichtermann for maintaining the nematode stock cultures, s, and to Justyna Frankowska-Łukawska and Ewa Znojek for perfect microtomy and ultramicrotomy assistance. Fungal cultures were kindly provided by Prof. Matthias Hahn, TU Kaiserslautern, Germany.

## Supporting information legends

**Figure S1: Cyst nematode infection assay in Col-0 and mutants of 10 selected candidate genes**.

**Figure S2: Genotyping of *hipp27* mutant lines**.

**Figure S3: RT-PCR analysis of *HIPP27* expression in Col-0 and *hipp27* mutant lines**.

**Figure S4: Phenotyping of Col-0 and *hipp27* mutant line grown under different growth conditions**.

